# Advances in abscission signaling

**DOI:** 10.1101/122168

**Authors:** O. Rahul Patharkar, John C. Walker

## Abstract

Abscission is a process in plants for shedding unwanted organs such as leaves, flowers, fruits, or floral organs. Shedding of leaves in the fall is the most visually obvious display of abscission in nature. The very shape plants take is forged by the processes of growth and abscission. Mankind manipulates abscission in modern agriculture to do things like prevent pre-harvest fruit drop prior to mechanical harvesting in fruit orchards. Abscission occurs specifically at abscission zones that are laid down as the organ that will one day abscise is developed. A sophisticated signaling network initiates abscission when it is time to shed the unwanted organ. In this article, we review recent advances in understanding the signaling mechanisms that activate abscission. Physiological advances and roles for hormones in abscission are also addressed. Finally, we discuss current avenues for basic abscission research and potentially lucrative future directions for its application to modern agriculture.

## Introduction

Abscission in plants is the process of shedding unwanted organs. The most recognized abscission event occurs in the northern hemisphere when deciduous plants drop their leaves before winter. However, plants can abscise leaves, branches, flowers, floral organs (petals, sepals, stamen), fruits, and seed pods. In general, plants have a flexible ordered design wherein size, number of organs, and precise shape are not strictly set. If abscission did not occur, plants would look very different. For example, if deciduous trees did not abscise their leaves before winter, they would retain remnants of the previous year’s leaves all over themselves. While some of these remnants may break off, the dead leaves that remain would eventually decay and invite disease as well as shield some light from living leaves. In essence, the ordered design we see in plants is shaped by both growth and abscission. It is easy to imagine how reproduction in many plants could be adversely impacted if abscission did not exist. If fruits with seeds were to never fall to the ground, the seeds would not touch the soil without aid of outside forces, like animals or harsh weather.

Abscission can be triggered by developmental cues like fruit ripening or fertilization. Flower petals falling off after fertilization is well characterized in Arabidopsis. The environment can also prompt abscission. Photoperiod and cooler temperatures trigger leaf abscission in the fall. By losing leaves in the fall, plants save on energy needed to keep them alive. Also, trees without leaves have less surface area to catch snow and ice, which reduces the risk of branches breaking under excess weight. Plants also cut their transpirational load during drought by abscising leaves. Many plants, such as bean, shed entire flowers when exposed to drought. Without adequate water it would be hard to set seeds, so this adaptation prevents plants from wasting energy starting seed production that they would not be able to finish. Insect feeding also triggers abscission in plants as a protective strategy (Faeth *et al*., 1981). Leaves can also be shed as a response to bacterial disease. Leaf shedding as a defense response leaves the pathogen feeding on the fallen leaf, giving the plant time to mount a more comprehensive preventative defense response (Faeth *et al*., 1981). In short, abscission is used by sessile plants to regulate their morphology to suit the environment in which they live.

Modern agriculture manipulates the abscission process to its advantage. For example, synthetic auxins and ethylene blockers, which partially block abscission, are sprayed on *Citrus* and apple trees about a month before harvest (Anthony and Coggins Jr., 1999; Yuan and Carbaugh, 2007). This practice prevents fruits from dropping to the ground before mechanical harvesters can collect them. Tomatoes used in the canning industry are bred with the *jointless* mutation, which results in plants with no pedicel abscission zone (Mao *et al*., 2000). When *jointless* tomatoes are picked, they leave their calyx and stem behind on the plant. This results in less puncture damage to other tomatoes when they are placed together in a harvesting bag (Zahara and Scheuerman, 1988). The future looks bright for further agricultural improvements resulting from altering abscission. For instance, many crop plants are overly sensitive to periods of mild drought common in the agricultural setting. It may be possible to increase yield in beans by preventing flowers from abscising in response to mild drought conditions (Pandey *et al*., 1984).

Abscission occurs specifically at a specialized region of cells called the abscission zone (AZ). AZs are laid down early in development and often have a band-like appearance. The cells in an AZ are smaller than the surrounding cells and have a densely packed cytoplasm. Once abscission is triggered, AZ cells expand and the middle lamella (the pectin layer that glues two cells together) is dissolved via hydrolytic enzymes, allowing cell separation. After abscission has occurred, a new, protective epidermal layer is laid down over the abscission “scar.”

## The abscission signaling network in Arabidopsis

For years Arabidopsis floral organ abscission has served as the premiere model for understanding abscission at the molecular genetic level. Forward and reverse genetic approaches have revealed a number of components that are necessary for abscission and recently biochemistry has led to some mechanistic insights. Abscission can be divided into 4 phases to simplify its explanation. First, abscission zones must develop. Second, abscission signaling is activated. Third, an enzymatic hydrolysis of the abscission zone’s middle lamella takes place and AZ cells begin to enlarge. Finally, the abscission scar further differentiates and seals itself (Kim, 2014). The hydrolysis phase has been thoroughly reviewed and will not be discussed in detail here (Niederhuth *et al*., 2013; Kim, 2014). A model of the physiological phases of abscission is shown in Figure 1.

**Figure 1.**
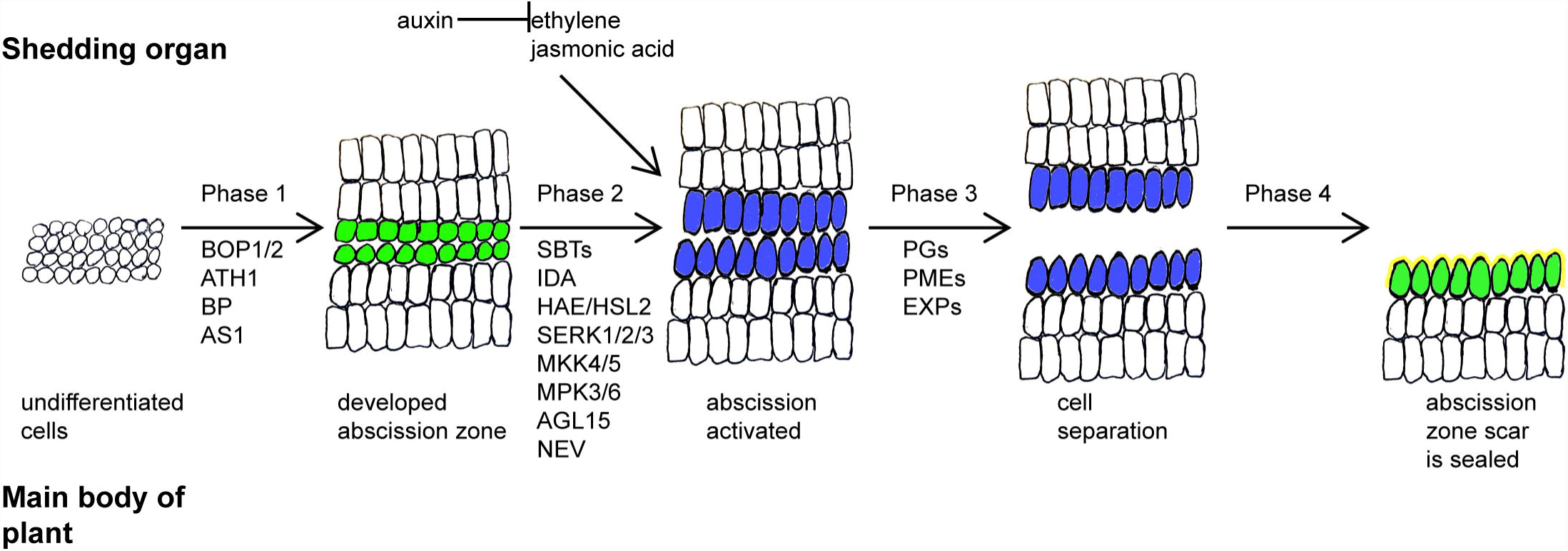
Physiological model of abscission. In phase 1, abscission zones develop. BOP1/2, ATH1, BP, and AS1 are required for abscission zone development. In phase 2, the abscission signaling loop is activated and the indicated proteins are all required. The hormones ethylene and jasmonic acid positively regulate abscission, while auxin negatively regulates abscission by reducing the effect of ethylene. At this step, abscission zone cells begin to enlarge and the pH of their cytosol becomes alkaline. In phase 3, polygalacturonases (PGs), pectin methyltransferases (PMEs), and expansins (EXPs) cause cell separation. These same enzymes, particularly expansins, are likely involved in the cell expansion that occurs before cell separation. Finally, in phase 4, the abscission zone scar is sealed with a protective layer and the pH of the abscission zone cells returns to neutral. Abscission zone cells are depicted in color where green represents neutral cytosolic pH and blue represents alkaline pH. This figure was adapted and updated from (Kim, 2014).

AZ development begins very early in the development of the organ that will later be able to abscise. Some genetics is known about AZ development in Arabidopsis. The transcription factors *BLADE ON PETIOLE 1/2 (BOP1/2)* are redundantly necessary for its formation (McKim *et al*., 2008). ARABIDOPSIS THALIANA HOMEOBOX GENE1 (ATH1), a BELL-type transcription factor, is required for stamen AZ placement and development (Gómez-Mena and Sablowski, 2008). The homeodomain transcription factor BREVIPEDICELLUS (BP) prevents floral organ AZs from becoming too big (Wang *et al*., 2006). The myb transcription factor, *ASYMMETRIC LEAVES1 (AS1)*, establishes the positioning of floral organ AZs (Gubert *et al*., 2014). In tomato, an agamous-like MADS domain transcription factor, *JOINTLESS*, is necessary for formation of the pedicel AZ (Mao *et al*., 2000). While MADS domain transcription factors regulate abscission activation in Arabidopsis, they have not been shown to regulate AZ development in Arabidopsis (Fernandez *et al*., 2000; Chen *et al*., 2011; Patharkar and Walker, 2015; Patharkar *et al*., 2016).

There is a broad understanding of the mechanisms that result in the activation of abscission signaling in Arabidopsis. Signaling events in the abscission activation phase initiate the expression of a mixture of cell wall modifying enzymes that dissolve the pectin-rich middle lamella of the AZ. No known Arabidopsis mutants abscise but then fail to further differentiate their AZs after abscission. However, it is likely that signaling components of the abscission activation phase play a role in the final differentiation of the AZ scar. For example, overexpression of *INFLORESCENCE DEFICIENT IN ABSCISSION (IDA)*, a gene needed for abscission activation, results in over-differentiation of the AZ scar (Stenvik *et al*., 2006).

## The abscission receptor complex and perception of processed IDA peptide

Abscission activation is regulated by two receptor-like protein kinases, *HAESA* and *HAESA-like 2 (HAE/HSL2)*, which are redundantly required for Arabidopsis to shed its petals, sepals, and stamen (Jinn *et al*., 2000; Cho *et al*., 2008; Stenvik *et al*., 2008). A peptide released from IDA by subtilisin-like serine proteinase processing is also required for abscission (Butenko *et al*., 2003; Schardon *et al*., 2016). Recent work indicates that HAE does not work alone to perceive IDA; rather, HAE works together with SOMATIC EMBRYOGENESIS RECEPTOR-LIKE KINASE 1/2/3/4 (SERK1/2/3/4) (Meng *et al*., 2016; Santiago *et al*., 2016). *SERK3* is also known as *BRI1-ASSOCIATED RECEPTOR KINASE 1 (BAK1)*. Importantly, BAK1 is also the co-receptor for both BRASSINOSTEROID INSENSITIVE 1 (BRI1) and FLAGELLIN-SENSITIVE 2 (FLS2) (Chinchilla *et al*., 2007; Nam and Li, 2002). BRI1 is the receptor that perceives brassinosteroid hormones that are responsible for cell expansion and elongation (He *et al*., 2000). FLS2 is the receptor that perceives a portion of bacterial flagellin and initiates a defense response (Gómez-Gómez and Boller, 2000). IDA stabilizes a protein complex between HAE/HSL2 and SERK 1/2/3/4 in Arabidopsis mesophyll protoplasts and in *Nicotiana benthamiana* epidermal cells (Meng *et al*., 2016). Specifically, the extracellular domains of HAE/HSL2 and SERK 1/2/3 interact with the IDA peptide (Meng *et al*., 2016; Santiago *et al*., 2016). Notably, an IDA peptide-HAE-SERK complex has only been shown to exist *in vitro* as well as in Arabidopsis mesophyll protoplast and in *Nicotiana benthamiana* overexpression systems (Meng *et al*., 2016; Santiago *et al*., 2016); this complex has yet to be identified in AZs. It will be interesting to see if new intricacies of the ligand-receptor complex can be uncovered in AZs.

Recent developments have shed new light on how IDA is processed into a biologically active form and shown the mechanism of IDA’s proteolytic processing into a biologically active peptide. A recent study overcame functional redundancy of subtilisin-like proteinases (SBTs) by expressing tissue-specific proteinase inhibitors to show that SBTs are both required for maturation of the IDA peptide and floral organ abscission (Schardon *et al*., 2016). Furthermore, the study revealed that the mature and highly bioactive IDA peptide is a 14 mer of sequence GVPIPPSAPSKRHN. This mature IDA peptide is at least 10 times more bioactive than the previously proposed PIPP or extended PIPP peptide previously proposed based on sequence conservation (Stenvik *et al*., 2008; Schardon *et al*., 2016). The exact contribution of specific SBTs to IDA cleavage and the extent of redundancy are not clear at the moment. SBTs 4.12, 4.13, and 5.2 are all proposed to contribute to IDA processing based on their expression patterns and *in vitro* activity (Schardon *et al*., 2016). Interestingly, the *in vitro*, mesophyll protoplast, and *Nicotiana benthamiana* over-expression studies that were used to show that IDA peptide can induce a complex between HAE and SERKs require a version of the peptide where the central proline is modified to hydroxyproline (Meng *et al*., 2016; Santiago *et al*., 2016). In contrast, studies involving AZs from Arabidopsis require no such modification for complementation of abscission defects of *ida* mutants or plants blocked in SBT activity (Stenvik *et al*., 2008; Schardon *et al*., 2016). This difference between findings in AZs and other non-AZ systems represents an opportunity to further refine our understanding of the abscission signaling mechanism.

## Amplification of the abscission signal by a MAPK cascade and a positive feedback loop

Downstream of the HAE receptor complex lies a MITOGEN-ACTIVATED PROTEIN KINASE (MAPK) cascade consisting of MKK4/5 and MPK3/6. Knockouts/knockdowns of the MAPK cascade fail to abscise their floral organs. Conversely, expression of constitutively active versions of the MKK4/5 are able to restore abscission in *hae hsl2* double mutants, indicating that the MAPK cascade is epistatic to the HAE receptor complex (Cho *et al*., 2008). At the moment, it is not clear which MAP triple kinase functions in the abscission pathway, nor is it clear whether there are other intermediates between the HAE receptor complex and the MAPK cascade. A striking discovery revealed that knockdown of MKK4/5 results in less than 20% of normal *HAE* expression as abscission is activated in floral organ abscission zones (Patharkar and Walker, 2015). This result is surprising since *HAE* was thought to be genetically upstream from the MAPK cascade (Cho *et al*., 2008). Over-expression of the MADS domain transcription factor, *AGAMOUS-LIKE 15 (AGL15)*, blocks abscission but does not alter AZ development, suggesting AGL15 is a negative regulator of abscission (Fernandez *et al*., 2000). A study in floral receptacles revealed that under native protein levels, AGL15 binds the *HAE* promoter keeping *HAE* from being expressed prior to abscission being activated. Furthermore, once the abscission signaling pathway is activated, the MAPK cascade phosphorylates AGL15 on serine 231 and 257 and de-represses *HAE* expression. Newly synthesized HAE completes a positive feedback loop once it takes its place in the plasma membrane. The positive feedback loop and the MAPK cascade both serve to greatly amplify the starting signal to abscise, which explains how *HAE* expression is increased 27-fold during the process of floral organ abscission. While AGL15 appears to be a major transcription factor regulating abscission, it certainly cannot be the only one. For example, *agl15 agl18* double mutants abscise statistically earlier than wild type, suggesting that AGL15’s sister protein, AGL18, plays a partially redundant role with AGL15 (Patharkar and Walker, 2015; Patharkar *et al*., 2016). Other transcription factors that block abscission when over-expressed have previously been reviewed (Niederhuth *et al*., 2013). Currently, it is not entirely clear how these other transcription factors fit into the aforementioned positive feedback loop.

Proper abscission requires an ADP-ribosylation factor GTPase-activating protein, NEVERSHED (NEV). Mutations in NEV alter the Golgi structure and change the location of the trans Golgi network. *nev* mutants also over accumulate paramural vesicles, which are thought to move HAE and other proteins to the plasma membrane (Liljegren *et al*., 2009). However, this story is anything but simple. Mutations in three different secondary genes can partially restore vesicle trafficking and restore abscission in *nev* mutants. The first of these three secondary genes is *EVERSHED (EVR)*, a receptor-like protein kinase that is also known as *SUPPRESSOR OF BIR1 1 (SOBIR1)* (Leslie *et al*., 2010). Mutations in *BAK1-INTERACTING RECEPTOR-LIKE KINASE 1 (BIR1)* have a constitutive pathogen response that can be suppressed by secondary mutations in *EVR/SOBIR1* (Gao *et al*., 2009). Secondary mutations in *SERK1* can also suppress *nev* phenotypes (Lewis *et al*., 2010). From a molecular mechanistic standpoint, it is not clear how mutating the co-receptor of HAE could restore abscission in *nev* mutants. The triple mutant *serk1 serk2 bak1* actually has a mild floral organ abscission defect (Meng *et al*., 2016). Finally, secondary mutations in *CAST AWAY (CST)*, a receptor-like cytoplasmic kinase, can also suppress phenotypes of *nev* mutants. CST physically interacts with HAE and EVR in Arabidopsis mesophyll protoplasts (Burr *et al*., 2011). Secondary mutations in *EVR, SERK1*, and CST all restore abscission in *nev* mutants, but the final AZ scar in these plants is over differentiated. This observation suggests that players involved in the abscission activation phase also function in the final differentiation of the AZ scar. In addition to being shuttled by vesicles, HAE passes through an error checking mechanism in the endoplasmic reticulum as well. The endoplasmic reticulum-associated degradation system (ERAD) ensures HAE is free from defects (Baer *et al*., 2016). When the ERAD system is defective, alleles of *HAE* that generate a partially functional protein can still make it to the plasma membrane and transduce the abscission signal. A model of the abscission activation signaling pathway is shown in Figure 2 and Table 1.

**Figure 2.**
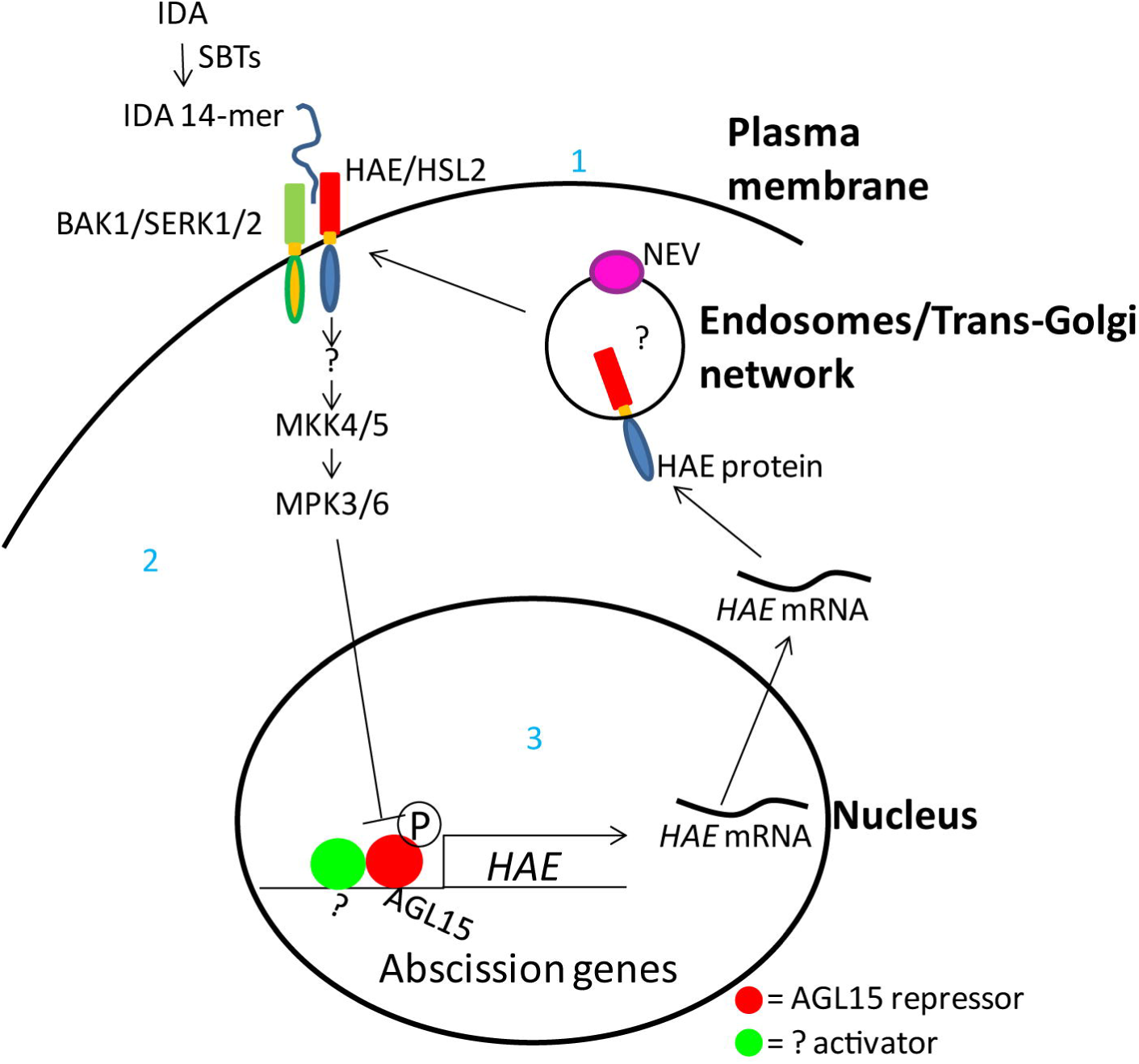
Model of abscission activation signaling pathway. IDA is cleaved by SBTs into a 14-mer peptide that binds HAE and its co-receptor that can be SERK1/2/3 (SERK3 is also known as BAK1). HAE then, through unknown means, activates a MAPK cascade consisting of MKK4/5 and MPK3/6. MPK3/6 then phosphorylate AGL15, which de-represses *HAE* transcription. Newly synthesized HAE is then shuttled back to the plasma membrane in endosome vesicles with the assistance of NEV, completing a positive feedback loop. Blue colored numbers indicate additional components that are located at the plasma membrane (1), cytoplasm (2), and nucleus (3) that cannot be precisely placed on the diagram. The names of these additional proteins are listed in Table 1 and were previously reviewed in detail (Niederhuth *et al*., 2013).

**Table 1.** Additional abscission signaling components not precisely placed on Figure 2.

| Cellular location | Protein | Reference |
| --- | --- | --- |
| 1 (plasma membrane) | CST | (Burr *et al.*, 2011) |
| 1 (plasma membrane) | EVR | (Burr *et al.*, 2011) |
| 1 (plasma membrane) | ETR1 | (Patterson and Bleecker, 2004) |
| 2 (cytoplasm) | EIN2 | (Patterson and Bleecker, 2004) |
| 3 (nucleus) | AGL18 | (Adamczyk *et al.*, 2007) |
| 3 (nucleus) | FYF | (Chen *et al.*, 2011) |
| 3 (nucleus) | COI1 | (Kim *et al.*, 2013) |
| 3 (nucleus) | BP | (Shi *et al.*, 2011) |
| 3 (nucleus) | KNOTTED-LIKE FROM ARABIDOPSIS<br>THALIANA 2/6 | (Shi *et al.*, 2011) |
| 3 (nucleus) | ZINC FINGER PROTEIN 2 | (Cai and Lashbrook, 2008) |
| 3 (nucleus) | DNA BINDING WITH ONE FINGER 4.7 | (Wei *et al.*, 2010) |

## Physiology, hormones, and the big picture of abscission

At first glance, the molecular mechanisms regulating abscission, that are described above, seem relatively straightforward and logical. However, their depiction is overly simplified since there are several points that are only partially congruent with physiological observations. Therefore, a great many opportunities exist to connect our understanding of molecular mechanisms regulating abscission with the actual physiological changes that occur in AZs during abscission. In our opinion a glaring issue that could be clarified is what exactly does IDA do at the physiological level? Recent literature refers to IDA peptide as a hormone, which it is likely to be (Santiago *et al*., 2016; Schardon *et al*., 2016). However, hormones are typically defined as molecules produced in one tissue that exert an effect in another tissue. Currently, no effort has been made to determine where the IDA peptide acting on a given AZ cell is coming from: the same cell, the immediately adjacent cells, or from more distant cells? Also, why is there a peptide signal at all? Does it help synchronize the abscission process? As drawn in model diagrams, all the molecular components necessary for signaling abscission are produced in each individual cell. *In vitro* experiments show that IDA peptide in agar plates can enter the pedicel of detached flowers and complement *ida* mutants (Stenvik *et al*., 2008). In short, there could be many future breakthroughs at the interface of molecular signaling and the physiology of complex AZ tissue. An increased understanding of AZ tissue will ultimately push mankind’s understanding of cell to cell communication to a new level.

A number of more classical plant hormones exert influence over abscission. Ethylene is broadly necessary for abscission in Arabidopsis and crop plants. Arabidopsis mutants defective in ethylene perception, *ethylene response 1 (etr1)*, and ethylene signaling, *ethylene insensitive 2 (ein2)*, have delayed floral organ abscission (Patterson and Bleecker, 2004). Additionally, the MADS domain transcription factor *FOREVER YOUNG FLOWER (FYF)* has been proposed to work at the level of ethylene signaling. Over-expression of *FYF* results in delayed abscission (Chen *et al*., 2015). Auxin is generally thought to negatively regulate abscission by making tissue insensitive to ethylene (Sexton and Roberts, 1982). As mentioned above, a cocktail of ethylene blocker (aminoethoxyvinylglycine HCl) and synthetic auxin (2,4-Dichlorophenoxyacetic acid) are used to prevent pre-harvest fruit drop in *Citrus* and apple tree (Anthony and Coggins Jr., 1999; Yuan and Carbaugh, 2007). Jasmonic acid positively regulates floral organ abscission in Arabidopsis. Mutations in the jasmonic acid receptor, *coronatine insensitive 1 (coi1)*, result in delayed floral organ abscission in Arabidopsis (Kim *et al*., 2013). Hormones other than ethylene, auxin, or jasmonic acid are also likely to influence abscission. Salicylic acid may also regulate abscission. The genes encoding enzymes necessary for salicylic acid synthesis, *ISOCHORISMATE SYNTHASE 1/2*, are transcriptionally increased during the process of floral organ abscission (Cai and Lashbrook, 2008). Salicylic acid has a well-established role in regulating senescence, and both floral organs and cauline leaves appear to senesce before they abscise (Guiboileau *et al*., 2010; Patharkar and Walker, 2015, 2016).

Many interesting yet relatively unexplained physiological events occur in AZs once abscission is activated. As abscission advances, the cytosol of AZ cells becomes more alkaline. Treatments that slow abscission, like ethylene blockers, prevent this cytosolic alkalization (Sundaresan *et al*., 2015). Additionally, cytosols of ethylene insensitive mutants in Arabidopsis do not alkalize, while mutants with overly active ethylene signaling are already alkalized before abscission begins. *ida* and *nev* mutants also fail to alkalize the cytosol of their AZ cells. The cytosolic alkalization of AZ cells, associated with abscission, has been shown to occur in Arabidopsis, tomato, and wild rocket (Sundaresan *et al*., 2015). The reason for the pH change of AZ cell’s cytosol is currently a mystery. One hypothesis is that alkaline pH may be optimal for some abscission enzymes.

Abscission zone cells also enlarge as abscission occurs. Arabidopsis cauline leaf AZ cells can clearly be seen to begin enlarging slightly prior to abscission. Thus, AZ middle lamella hydrolyses and AZ cell enlargement overlap in timing. In drought triggered-abscission, the final size of AZ cells that have a sealed scar are not noticeably larger than AZ cells at the moment of first cell separation (Patharkar and Walker, 2016). Previous reviews suggest that floral organ AZ cell enlargement only happens after cell separation is complete (Kim, 2014). This notion is likely due to the fact that floral organ AZ cells cannot be visualized nondestructively prior to abscission because sepals and petals cover the AZs. After learning cauline leaf AZ cells were expanding and separating simultaneously, we looked closely at floral organ AZs as cell separation was just beginning. When we removed loosely attached sepals and petals (i.e., when abscission is just beginning but not complete), we observed already enlarged AZ cells. It should be noted that early in the 20^th^ century scientists believed mechanical shearing force from AZ cell enlargement was the primary driving force for abscission (Fitting, 1911; Sexton and Roberts, 1982). Currently, no mutants of Arabidopsis abscise but fail to enlarge their AZ cells suggesting that AZ enlargement is necessary for abscission in Arabidopsis.

## Leaf abscission and abscission in non-model plants

The most detailed explanation of abscission signaling has come from studying Arabidopsis floral organ abscission. The Arabidopsis floral organ system has been used to work out a number of molecular mechanisms regulating abscission signaling (described above) that would have been more difficult to solve in less genetically tractable systems. Floral organ abscission occurs once fertilization occurs. Since Arabidopsis is self-pollinating, abscission basically occurs in a developmentally timed manner. Recently, it has become clear that cauline leaf abscission triggered by drought requires the same core abscission signaling mechanism as floral organ abscission (Patharkar and Walker, 2016). *IDA, HAE/HSL2, MKK4/5*, and NEV are all required for drought-triggered leaf abscission to occur. This is an interesting finding because cauline leaf abscission is not set on a developmental clock but rather occurs conditionally. Drought to the point of wilting activates HAE expression, and then, once plants are re-watered, leaf abscission occurs (Patharkar and Walker, 2016). HAE expression is also activated in the vestigial pedicel abscission zone in Arabidopsis prior to partial abscission. This observation suggests that fruit abscission may also utilize the same signaling pathway as leaves and floral organs (Patharkar and Walker, 2016). The cauline leaf abscission system in Arabidopsis has two distinct features from the floral organ abscission system that will likely aid researchers in further unraveling the process of abscission. First the cauline leaf AZ can be non-destructively observed from development through abscission because no tissue obscures its view. Second, abscission is not triggered by a developmental stage, rather it is triggered by environmental conditions. This allows separation of abscission events from developmental events. Currently, drought in known to trigger cauline leaf abscission, however, other environmental stimuli may also initiate leaf abscission (Patharkar and Walker, 2016).

How conserved is the Arabidopsis abscission signaling module in other species? Recent research indicates that the Arabidopsis abscission module likely extends far past Arabidopsis. A phylogenetic study showed that IDA is conserved in all flowering plants (Stø *et al*., 2015). *HAE* homologs are up-regulated in Poplar leaf AZs that are abscising due to shading (Jin *et al*., 2015). *Citrus IDA3* can complement abscission deficiency of Arabidopsis *ida* mutants (Estornell *et al*., 2015). Taken together, these findings are strong evidence that the Arabidopsis abscission signaling module works in other and distantly related species.

## Future research

In our opinion, two areas of basic abscission research stand out as particularly likely to pay big dividends in the future. First, a literature cross reference analysis can provide a number of easy to test leads to extend the existing abscission signaling pathway. Since abscission components overlap with defense and brassinosteroid fields, and drought-triggered abscission has been characterized, mining those fields for connections relevant to abscission could be fruitful. For example, one could look at the defense field and see that BIR1 interacts with BAK1 and that mutations in *EVR* can suppress the phenotypes of *bir1* mutants. The power of cross reference analysis will grow as more network hub components are added to the abscission pathway. The ability to work in both floral organ and cauline leaf abscission zones also gives researchers options. Of course, researchers will have to take care in interpreting non-AZ data and applying it to AZs. The AZ is a unique tissue that does not behave like other parts of the plant. For example, AGL15 over-expression represses HAE expression in AZs but not in mesophyll protoplasts (Patharkar *et al*., 2016).

A second promising area is research focused on precisely connecting molecular data to the physiology of abscission zones. At the moment, molecular signaling mechanisms have very vague outputs. For the most part, all that can be said for the output of the molecular pathway is that it triggers abscission. However, abscission can be broken down into a number of physiological events, like AZ cell enlargement, AZ cell cytosol alkalization, and middle lamella hydrolysis. How does the molecular pathway affect these different physiological aspects? It is unlikely that all AZ cells behave the same. Using modern physiological methods to break down the AZ into functional groups of cells and events will yield a tangible understanding of what actually happens at the cellular level to allow abscission.

Finally, there is also promise to use our basic knowledge of abscission signaling to do applied research that could benefit agriculture. For example, preventing soybeans from shedding flowers in response to mild drought conditions might result in more seed pods come harvest time. *ida* and *hae hsl2* mutants are abscission defective but have no seed yield penalty, making them excellent candidates for manipulation (Patharkar and Walker, 2016). Knocking out their homologs in soybean via CRISPR/Cas9 is doable and would allow for straightforward seed yield trials. Alternatively, yield trials could be conducted where chemicals that delay abscission are sprayed on plants prior to drought treatment. While there is no guarantee that the proposed manipulations to soybean would result in increased grain yield, the mechanism for grain yield improvement is simple and therefore straightforward to assay. Basically, more flowers on the plant could be more seed pods later on. In summary, there are more opportunities in basic and applied abscission research right now, than ever before.

## Acknowledgments

We would like to thank Catherine Espinoza, Melody Kroll, and Isaiah Taylor for reading and editing the manuscript.

